# Predicting the Emergence of SARS-CoV-2 Clades

**DOI:** 10.1101/2020.07.26.222117

**Authors:** Siddharth Jain, Xiongye Xiao, Paul Bogdan, Jehoshua Bruck

## Abstract

Evolution is a process of change where mutations in the viral RNA are selected based on their fitness for replication and survival. Given that current phylogenetic analysis of SARS-CoV-2 identifies new viral clades after they exhibit evolutionary selections, one wonders whether we can identify the viral selection and predict the emergence of new viral clades? Inspired by the Kolmogorov complexity concept, we propose a generative complexity (algorithmic) framework capable to analyze the viral RNA sequences by mapping the multiscale nucleotide dependencies onto a state machine, where states represent subsequences of nucleotides and state-transition probabilities encode the higher order interactions between these states. We apply computational learning and classification techniques to identify the active state-transitions and use those as features in clade classifiers to decipher the transient mutations (still evolving within a clade) and stable mutations (typical to a clade). As opposed to current analysis tools that rely on the edit distance between sequences and require sequence alignment, our method is computationally local, does not require sequence alignment and is robust to random errors (substitution, insertions and deletions). Relying on the GISAID viral sequence database, we demonstrate that our method can predict clade emergence, potentially aiding with the design of medications and vaccines.

## Introduction

The outbreak of coronavirus disease 19 (COVID-19), caused by the severe acute respiratory syndrome coronavirus 2 (SARS-CoV-2; previously known as 2019-nCoV)^1^, has become a major pandemic posing a serious threat to global public health. The rapid worldwide spread of COVID-19 accompanied by a viral mutation-driven evolution has demanded researchers to study its genetic diversity. To better understand the evolution of SARS-CoV-2, computational mathematical approaches are urgently needed to uncover the information contained in currently updated databases, characterize the viral evolution^2, 3^, and help at designing efficacious vaccines. Towards this aim, we propose a universal novel computational method that enables us to analyze the viral RNA sequences and characterize the viral evolution and selection.

Figure 1 (a) illustrates an initial viral RNA strain (root clade) undergoing evolution dependent on environmental and stochastic factors giving rise to *transient* mutations. Some of these mutations get selected based on their fitness for replication and survival. These selected mutations are *stable* and give rise to a new clade that can further evolve into new subclades. Figure 1 (b) shows the phylogeny of SARS-CoV-2 clades obtained from nextstrain^4^. One might ask if the emergence of future viral clades can be predicted. To answer this question, we propose a mathematical model mapping an RNA sequence onto a state machine, a directed weighted network where the nodes and edges represent the states (i.e., contiguous groups of *k* nucleotides) and the transition probabilities among states, respectively (see Figure 2 (c)-(d)). The state machine with transition probabilities is also known as a Markov chain. In order to predict the future *b* nucleotides from previous *k* contiguous nucleotides in an RNA sequence, we develop a Markov chain consisting of 4^*k*^ states transitioning into 4^*b*^ states (see Figure 2 (a)-(b)). After traversing the entire RNA sequence, the number of occurrences of each transition provides information about the corresponding transition probability and also represents the weight of the corresponding directed edge (i.e., a directed weighted network of the RNA sequence is generated with 4^*k*^ nodes (see Figure 2 (e)-(f))). Figure 2 (a) and (b) show a transition *acg→t* with model *k* = 3, *b* = 1. Figure 2 (c) and (d) show a transition *acgt→cg* with model *k* = 4, *b* = 2. This proposed method encodes the evolution of RNA sequences over time and the transition probabilities reveal important characteristics that cannot be directly captured from RNA sequences.

**Figure 1.**
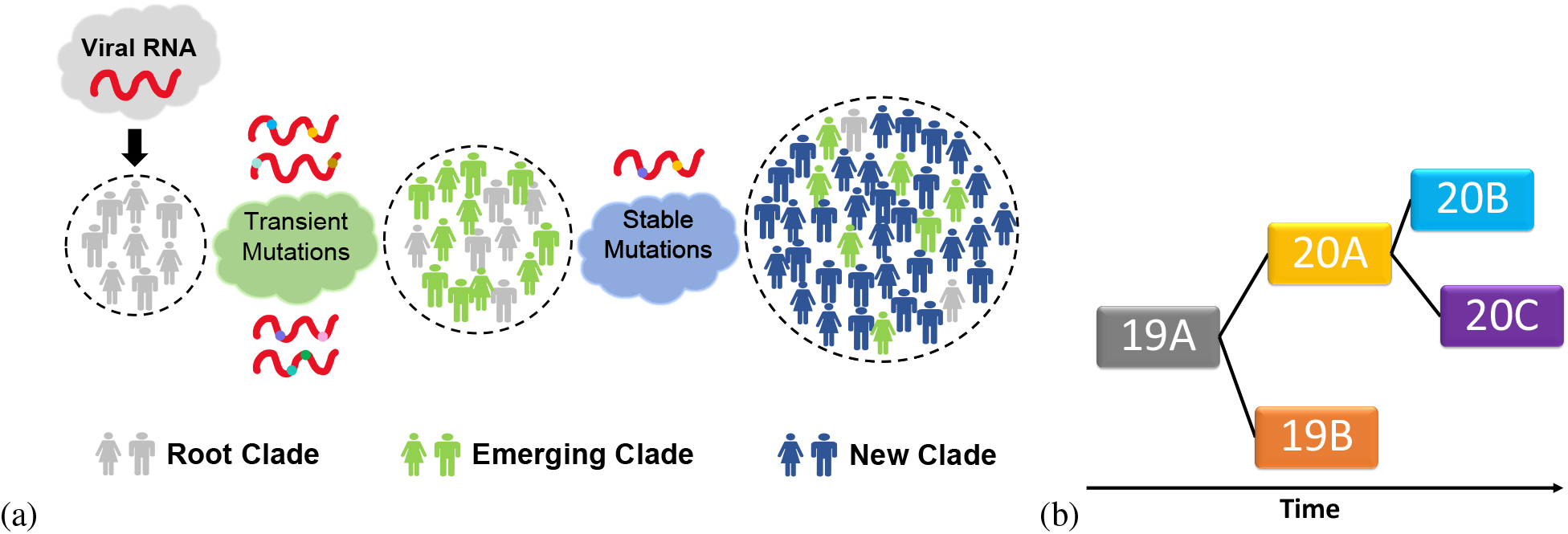
Evolution of a clade. (a) The initial viral RNA strain (root clade) undergoes evolution due to the environmental and stochastic factors to give rise to transient mutations which are then selected based on their fitness for replication and survival. The selected (stable) mutations give rise to a new viral strain (new clade) which can subsequently evolve into new subclades. The legends show people infected with different clades. (b) Phylogeny of SARS-CoV-2 clades obtained from nextstrain^4^.

**Figure 2.**
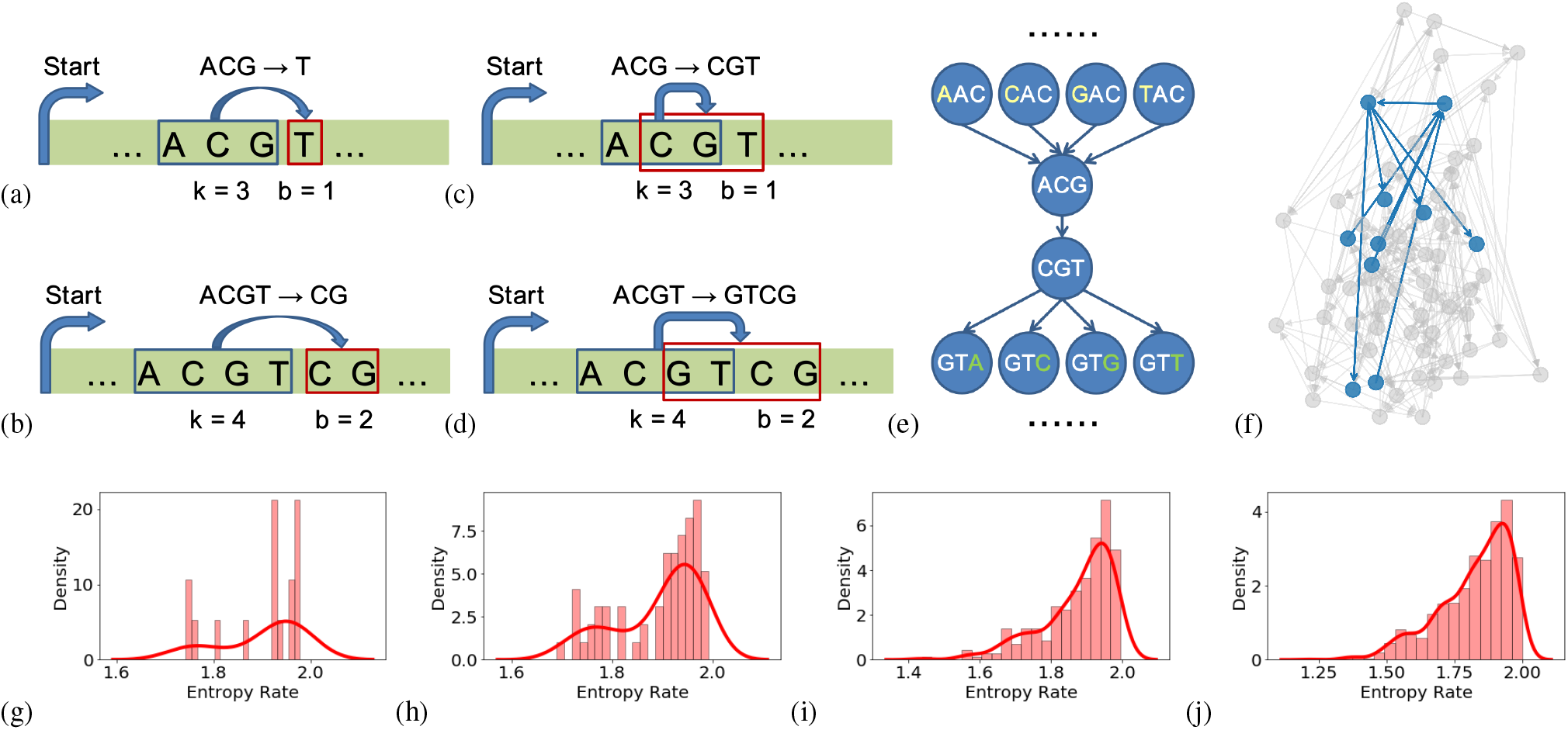
State machine model. The proposed state machine representation maps the RNA sequence onto a set of states and transition probabilities by identifying the start and next states of a varying number of nucleotides. For instance, a pair of start and next states of 3 nucleotides and 1 nucleotide is shown in (a) and 4 nucleotides and 2 nucleotides is shown in (b). In their corresponding directed weighted network representation, we consider each state to be a node and convert the transition probability into a weighted edge. Correspondingly, the model *k* = 3, *b* = 1 is shown in (c) and the model *k* = 4, *b* = 2 is shown in (d). A part of the directed graph showing the connection between states for the model *k* = 3, *b* = 1 is shown in (e), the overview of the whole directed graph is shown in (f) and the part shown in (e) is marked blue. (g)-(j) Entropy rate distribution for all the states in the state machine model for SARS-CoV-2 with *k* = 2, 3, 4 and 5 respectively with *b* = 1. For a state with *random* transitions, the entropy rate is maximum and is equal to 2.

## Results

### Higher-order interaction analysis on state machine representations of RNA sequences via entropy rate reveals the emergence of active transitions

Figure 2 (g)-(j) show the distribution of entropy rate for different states and for various state transition models. For the low order models (i.e., *k* = 2, 3), a high proportion of states have an entropy rate ranging between 1.8 - 2 bits/symbol, which means that the distribution of transitions to *a*, *c*, *g* and *t* is close to uniform. However, for the higher order models (i.e., *k* = 4, 5), there are about 15 - 20% of states with low entropy rate which means that those states are heavily biased towards one or two transitions taking up most of the probability mass. These transitions can be categorized as highly likely transitions. A few of those highly likely transitions that we found are *ccggg → t, cggat → g, gctcc → a, gtccc → t, gtctc → t* with the transition probability greater than 0.65. A list of transitions with high probability is provided in Supplementary Table S1. The mean entropy rate using the histogram given in Figure 2 (i) for *k* = 4, *b* = 1 is given by 1.895 *bits/nucleotide.* A high mean entropy rate shows that most of the transitions in the viral RNA sequence are uniformly random, a feature which we also see in Figure 2 (g)-(j).

### Classification of SARS-CoV-2 clades using the state machine model

To validate the robustness of the state machine model, we used the transition probability matrix obtained using the state machine model to compare sequences belonging to different clades - 19A, 19B, 20A, 20B, 20C identified by nextstrain^4^. Using 2700 samples, Figure 3 shows that our model is able to accurately compare all the clades. Figure 3 (a) shows the mean validation accuracy of pairwise classifiers (see Methods) built to compare all pairs of clades using transition probabilities as features. We see in Figure 3 (a) that the mean validation accuracy ranges between 99−100%. We also built a multiclassifier (see Methods) to compare all the clades together and show in Figures 3 (b)-(f) the mean classification probabilities greater than 0.9 for all the clades. This means that in the clade identification task, the multiclassifier not only identifies the correct clade but also with a very high probability. In other words, these pairwise and multiclassifiers can be used to accurately identify the clades. The advantage of our method of clade identification is that it doesn’t require *alignment* of sequences as the transition probability matrix calculated using the state machine model is robust to insertion, deletion and substitution mutations (see Supplementary Figures S3 and S4). *k* = 4, *b* = 1 is used to build pairwise classifier in Figure 3 (a) and *k* = 5, *b* = 1 is used to build the multiclassifiers in Figures 3 (b)-(e).

**Figure 3.**
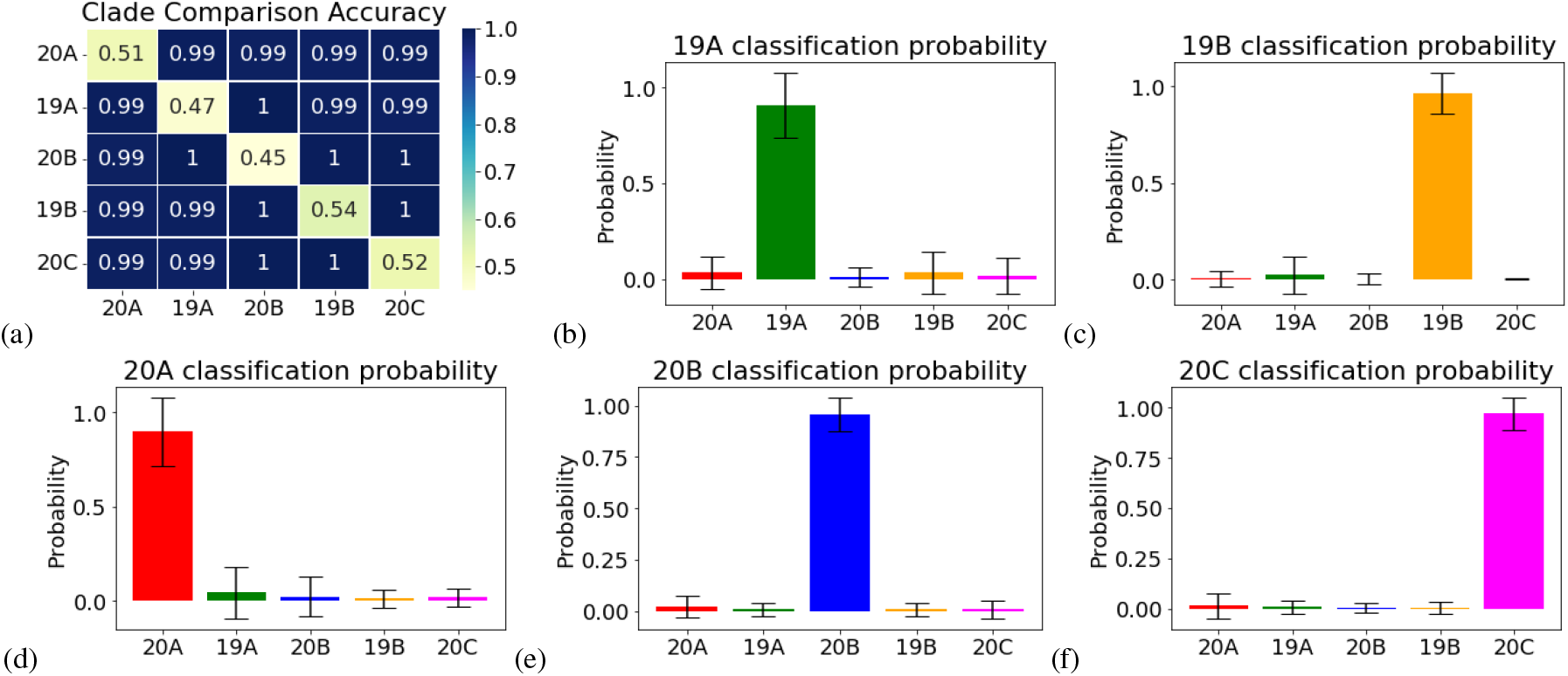
Clade classification using state machine model. The transition probability matrix associated with a state machine model with *k* = 4 and *b* = 1 can accurately distinguish the clades for SARS-CoV-2 identified by nextstrain^4^ as demonstrated by the high mean classification probabilities of built multiclassifier (greater than 0.9) for 19A (b), 19B (c), 20A (d), 20B (e), and 20C (f) clades. Figure 2(a) shows that the mean classification accuracy ranges between 99% and 100% for the pairwise classifiers built between different clade pairs. This state machine based clade classification is robust to insertion, deletion and substitution mutations and does not require sequence alignment.

### Mutation Detection using the state machine model

The mutations differentiating clades 19A, 19B, 20A, 20B, 20C are given in Supplementary Table S2. We show in Figure 4, that the state machine model can detect these mutations without *requiring alignment* of sequences. In this algorithm, we divide each viral RNA sequence into non-overlapping continuous regions of length *N* (i.e., ≈ 30000*/N* regions for SARS-CoV-2 virus). We construct the transition probability matrix for each of these regions and then build pairwise classifiers (see Methods) for each of these regions between the two clades to be compared. The regions with high validation accuracy point to areas where the transition probability matrices are distinguishable because of the presence of mutations. Therefore, the mutations are detected in the regions with high validation accuracy. In Figure 4 (a)-(j), we show the detected mutations (spikes with high accuracy) when different clades are compared with region length 1000. As can be seen in Figure 4 (a), we detect both A23403G (D614G, popularly known as spike protein mutation^5, 6^) and C14408T (P314L) mutations differentiating clade 20A with 19A. In fact, we also identify other mutations that haven’t been included on nextstrain in the definition of these clades. For example a mutation C241T in region 1 in Figure 4 (a)-(f) is also present in clade 20A, 20B and 20C when compared with clade 19A and 19B. We also notice another mutation C3037T in region 4 (3000 4000) when comparing clades 20A, 20B, 20C with 19A and 19B. Note that C3037T is sometimes also observed in region 3 (2000 3000) because of deletions and our method is able to detect its presence there. Note that in these curves, when accuracy is not close to 1 but in the range 0.7 - 0.8, it could either signify weaker detection or point to mutations that exist partially in a certain clade in that region, meaning they are present in a fraction of sequences of a given clade in that region. However, an accuracy close to 0.5 means no mutation is detected in that region.

**Figure 4.**
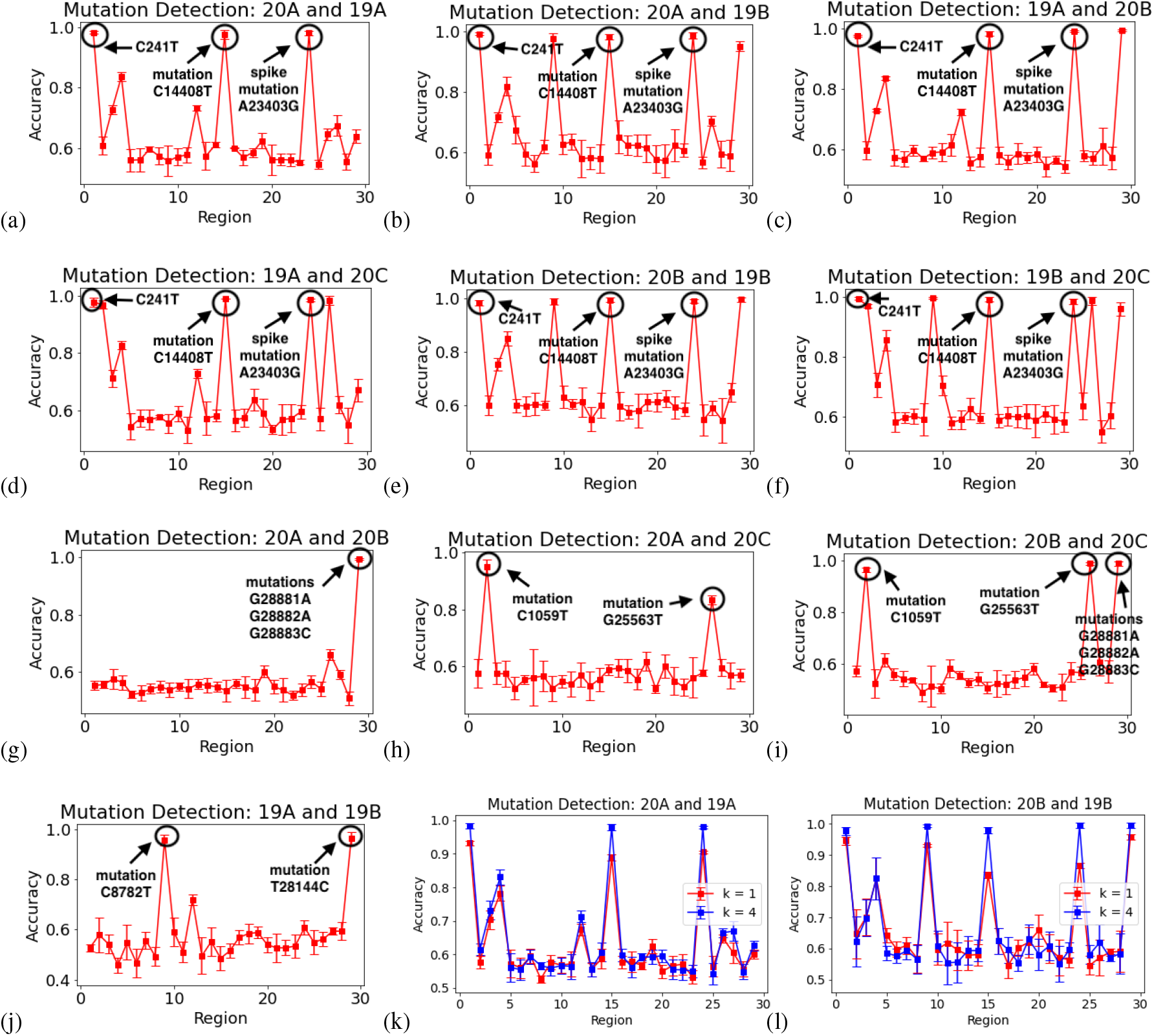
Mutation detection using the state machine model. Here, we show how the state machine model can be used to detect the mutations differentiating various clades without the need for sequence alignment. In this method, we divide each SARS-CoV-2 RNA sequence into 29 regions of length 1000 each and calculate the classification accuracy for the model comparing the transition probability matrix calculated for each 1000 length region. The regions for which the classification accuracy is high are the spots where the transition probability matrices are different due to mutations occurring in that region. For example, an accuracy of 1 in region 24 (23000 – 24000) in (a), (b), (c), (d), (e) and (f) indicates the detection of spike protein mutation (A23403G or D614G). In fact, a spike in region 1 (0 1000) indicates the mutation C241T in (a)-(f) which also appears in clades 20A,B,C when compared to clades 19A, B. We note that this mutation was not used in nextstrain to define clade 20A, B and C, but it is strongly correlated with C14408T and A23403G mutations. Further, in other plots, we also annotate mutations used by nextstrain to define clades 20B, 20C, 19B which were also detected using our state machine model. The state machine model used here has parameters *k* = 4 and *b* = 1. (k)-(l) Better detection accuracy is observed for higher order model with *k* = 4 compared to *k* = 1 with *b* = 1.

Figure 4 (k)-(l) show that the mutation detection is improved if we use *k* = 4 compared to *k* = 1 in the state machine model. Mutation detection with region length 500 and 100 are also provided (see Supplementary Figures S1 and S2). The mutation detection method is also robust to synthetically added noise in the form of substitution, insertion and deletion mutations (see Supplementary Figures S3 and S4).

### Evolution within clades and emergence of new clades

Using about 23000 complete and high coverage GISAID samples from clades 19A, 19B, 20A, 20B and 20C, we characterize *intra-clade* evolution over different months using the state machine model. In particular for a given clade, we compute the transition probability matrix for the samples and compare these matrices for earlier samples against matrices for samples in later months using machine learning classification (see Methods). We observe increasing mean validation accuracy of classification for clade 19A as the duration gap is increased as shown in Figure 5 (a). To analyze this finding, we conduct a deeper analysis to identify the regions that are highly active leading to this evolution observed within clade 19A. We divide the sequences into 10 continuous regions of length 3000 each and computed the transition probability matrices for all the samples in each region. For each region, we compare the transition probability matrices of samples in January against the samples in May for clade 19A using machine learning classification (see Methods). We find regions 3 (6000-9000), 4 (9000-12000), 5 (12000-15000), 8 (21000-24000) and 10 (27000-30000) with maximum validation accuracy as shown in Figure 5 (b) (left). Hence, by comparing transition probability matrices for 19A samples in the months of January and May in each region, *we identify regions of high mutation activity*. Interestingly, we also notice that the mutations characterizing clade 19B lie in regions 3, 4 and 10 (see Figure 5 (b) (top right)). It could be due to the evolving characteristics demonstrated here in regions 3, 4 and 10 that clade 19B emerged with mutations selected in those regions. Region 5 (12000-15000) and region 8 (21000-24000) are also found to have higher values for mean classification accuracy as shown in Figure 5 (b). Interestingly, two mutations (C14408T and A23403G) that define clade 20A are also found in regions 5 and 8, respectively. Further, another mutation is found in a fraction of 20A samples in region 4. It is possible that the emergence of clade 20A from 19A could be due to the mutation activity observed in these regions within clade 19A. Figures 5 (a)-(b) don’t include analysis for June samples due to lack of samples of clade 19A on GISAID in the month of June.

**Figure 5.**
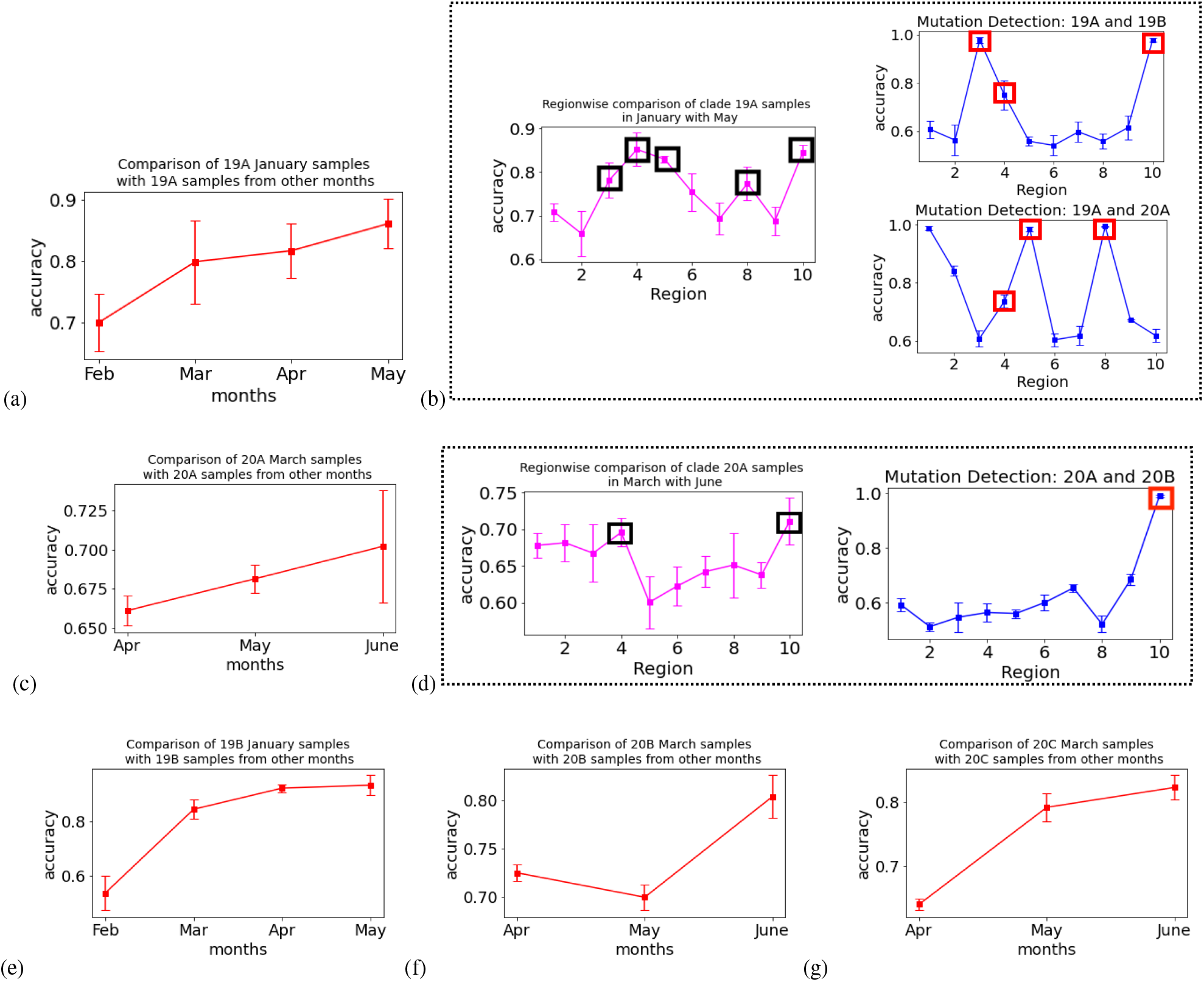
Evolution within clades and emergence of new clades. (a) Using state machine model, we observe that samples from clade 19A become more distinguishable (distant apart) from the earlier samples of clade 19A in January showing that the clade 19A is evolving and may further emerge into a new clade in the future. (b) The accuracy in May as observed in (a) is maximally contributed via regions 3 (6000 9000), 4 (9000 12000), 5 (12000 15000), 8 (21000-24000) and 10 (27000 30000). Interestingly, mutations differentiating clades 19A and 19B are also found in regions 3, 4 and 10 as highlighted in the right top figure. It could possibly mean that high mutation activity in regions 3, 4 and 10 within clade 19A led to the emergence of clade 19B defined by mutations in those regions. This may suggest that by finding the highly active areas within a clade, we can predict the regions of mutations that define a new clade. Further, two mutations (C14408T and A23403G) differentiating clade 20A from 19A lie in regions 5 and 8, respectively, which makes it plausible that these mutations were selected due to high activity of mutation in regions 5 and 8 within clade 19A. (c) Similar to clade 19A, clade 20A is also evolving with time. (d) Regions 4 and 10 possess the maximum classification accuracy within clade 20A and therefore are the most actively evolving areas within clade 20A. Interestingly the 3 mutations (G28881A, G28882A, G288883C) differentiating clade 20B from 20A are also observed in region 10 as demonstrated by the spike in the right figure. This makes it plausible that the emergence of these mutations in region 10 of clade 20B could be due to high mutation activity in region 10 within clade 20A. (e)-(g) Evolution within a clade is also observed for clades 19B, 20B and 20C, respectively.

Further, in Figure 5 (c), we demonstrate using the state machine model that clade 20A is also evolving with time. In Figure 5 (d), we identify regions 4 (9000-12000) and 10 (27000-30000) to be the most actively evolving within clade 20A on the basis of classification accuracy. Interestingly, clade 20B that emerged from clade 20A has 3 mutations G28881A, G28882A and G28883C that lie in region 10 (see Figure 5 (d) right). The emergence of these mutations in clade 20B could be due to the relatively higher evolution activity observed in region 10 within clade 20A.

We also observe intra-clade evolution over time for other clades - 19B, 20B and 20C in Figures 5 (e)-(g). We identify regions 6 (15000-18000), 7 (18000-21000), 8 (21000-24000) and 10 (27000-30000) to be active in the emergence of clade 19B (see Supplementary Figure S5 (a)). It is possible that in the future a new clade may emerge from 19B with mutations in regions 6, 7, 8 and 10. Region 10 is also one of the most active within clade 20B (supplementary Figure S5 (b)). Regions 1, 4 and 10 are the most active within clade 20C (supplementary Figure S5 (c)) indicating a possibility of the emergence of a new clade from 20C with mutations in these regions. As more samples of 19A, 19B, 20A, 20B, 20C from future months of July, August and further are collected, we hypothesize that active mutation regions that characterize the emergence of these clades can be identified more robustly using the state machine model and can be used to predict the emergence of new clades in the future for these clades. Figure 5 (e) doesn’t include the month of June due to the lack of samples of clade 19B on GISAID. Figures 5 (c), (d), (f), (g) don’t include the months of January and February due to the lack of samples of clades 20A, 20B and 20C on GISAID for those months.

## Discussion

By characterizing the higher order interactions among the RNA nucleotides within the viral RNA sequences, we provide new mathematical tools for investigating the evolution of viruses (i.e., SARS-CoV-2) and potentially enable the prediction of viral “dark matter”^3^. Our work can also enable the development of novel approaches to transform RNA or DNA sequences into new causal multiscale representations which preserve the distance between the sequences, store contextual information and are robust to insertion, deletion and substitution mutations. Further, this representation can be used to compare sequences without any need for alignment. The significance of our work can be two-fold: Firstly, the model can distinguish the virus mutations. Secondly, the proposed model can predict the virus mutation selection by investigating the evolution of the transition characteristics of the virus RNA sequences, which can help in developing sequence-based therapies, therapeutic agents and vaccines for viruses especially SARS-CoV-2^3, 7, 8^.

The state machine model we proposed can be viewed as a universal and interpretable tool to apply computational analysis on various DNA or RNA sequences including viral RNA sequences. Higher-order interactions among the RNA nucleotides could be computationally analyzed and used to characterize the RNA sequence, providing an approach to investigate viruses at the genetic level. As shown above, our model can be used to compare between different viral clades and detect mutations separating the clades without the need for alignment (Figures 3 and 4). We also capture the temporal characteristics of viral evolution. In particular, the model can be used to characterize the evolution within a clade (Figure 5 (a), (c), (e)-(g)) and in the identification of the actively mutating regions that could (Figure 5 (b), (d)) lead to the emergence of a new viral clade.

The viral RNA analysis for SARS-CoV-2 has been primarily focused on finding the phylogeny based on a first order model which considers the spelling mistakes or edit distance between the viral sequences^9–11^. A model based on the frequency of *k*-mers was proposed in the past for alignment free and global analysis of RNA and DNA sequences^12^. This model was recently used in^13^ to compare SARS-CoV-2 sequences with other viruses and earlier SARS-CoV-2 samples with the more recent ones. In this paper, we present a state machine based model for analyzing the viral RNA sequences. Similar to *k*-mer frequency based approach^12^, this model also doesn’t require alignment of sequences. In addition, it can be used to detect mutations, characterize the evolution and in the identification of the highly active regions within the virus where new viral mutations can emerge in the future.

Virologists can take our model and use it as a universal graph representation for all the viruses. The nodes in the graph represent the states and the edges are defined by the state transitions. The graph is universal as the graph structure is independent of the virus of interest. The weighted graph structure also represents a generative model for the virus of interest and can be used to define the distance between different viral types. This can aid us in identifying the associations between different viral types especially for the ones that haven’t been discovered/observed yet (i.e., it can help us understand and develop vaccines to deal with viruses that may appear in the future). Note that, our model is not limited to viral RNA and can be applied in principle to any RNA or DNA sequence.

It is critical for medical therapies and vaccine designs to not only understand the structure of the virus but also have knowledge of the viral dynamics. In particular, the emergence of likely mutations that can make the virus more infectious and lethal. Recently, it was claimed that the D614G mutation in the spike protein is causing the virus to be more infectious^5, 6^. In addition to the detection of spike protein mutation (A23403G or D614G) (Figure 4), we also find in Figure 5 (b) that region 8 (21000-24000) is an area of high evolution activity within clade 19A which could have led to the emergence or selection of the stable spike protein mutation that is found in clade 20A. Hence, by observing the mutation activity within a clade in different regions using the state machine model, it is possible to predict active areas in the SARS-CoV-2 RNA where critical mutations can emerge in the future. The location of these regions can then be used to identify viral proteins that may change their amino acid configuration in the future providing researchers valuable information required for robust vaccine design for COVID-19. Also, drug resistant mutants represent major obstacles to successful antiviral therapy. Thus, the insights into viral evolution are needed for the antiviral drug development^14^. Our model provides an approach to characterize the virus evolution and selection. Based on our model and RNA interference (RNAi)-based gene therapy^15^, we could focus on the direction of resistance mutations and use sequence-based therapy to prevent the selection of resistant mutations.

Further, our mathematical framework sheds light on the states in the virus which are less random (low entropy rate) compared to the ones which are random (high entropy rate). The low entropy states in the virus are more predictable as they have higher probabilities for certain transitions over the others giving us a tool to predict the most likely mutations.

There have been cases of COVID-19 in the past which have become severe in a short amount of time and lead to death in certain cases. Methods are needed to investigate the viral RNA sequences and predict their lethality before the appearance of severe symptoms. Further, these methods need to be robust in the sense that their false negative rate is as low as possible. The state machine model can also be used to detect infectious and lethal mutations in the SARS-CoV-2 sequence that might be leading to severe cases. However, there is an urgent need for comprehensive open datasets of viral RNA with symptomatic information to understand the connection between symptoms and the information contained in the viral RNA sequences in order to provide efficacious solutions to tackle the current and future pandemics.

## Methods

### Transition probability matrix and entropy rate

For a state machine model with parameters *k* and *b*, there are 4^*k*^ states and the empirical transition probability matrix *T* is estimated from the viral RNA sequence using the add-*β* estimator^16, 17^ given by

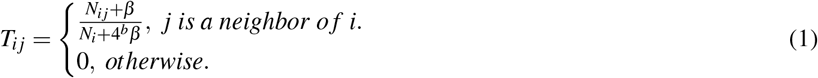

Here *i* ∈ {1, ⋯, 4^*k*^}. *j* is a neighbor of *i* if the last *k* − *b* symbols of *i* overlap with the first *k* − *b* symbols of *j* (see Figure 2). Therefore, there are 4^*b*^ neighboring *k*-mers of *i*. *N_ij_* denotes the number of occurences of *k*-mer *i* and its neighboring *k*-mer *j* together in the sequence (see Figure 2). *N_i_* denotes the number of occurences of *k*-mer *i* in the sequence. The entropy rate *E_i_* of state *i* is given by

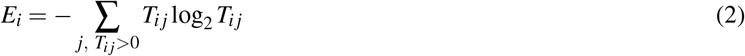

The empirical probability *p_i_* of state *i* is estimated using add-*γ* estimator^16, 17^ and is given by

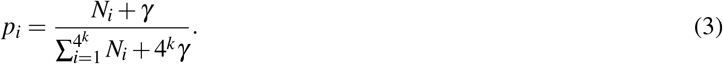

The empirical mean entropy rate *E* of the Markov chain is given by

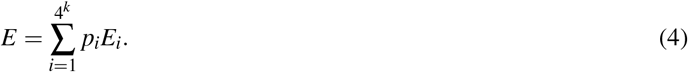

Note that, *β* = 0.5 and *γ* = 0.5 are used in all the experiments conducted in the paper.

### Machine learning classification

For a state machine model, the transition probability matrix is converted into a vector. We note that there can be at most 4^*k*+*b*^ non-zero entries in that vector as for any *k*-mer there are 4^*b*^ neighbors. We use this 4^*k*+*b*^ dimensional vector as the feature vector for building machine learning classifiers. In all the classification tasks, we have 2 stages. In the first stage, we perform feature extraction by building a learning model using XGBoost^18, 19^ (XGBoost package in python is used in the implementation) method with maximum depth = 1. In the next stage, we use the top 10 features identified in stage 1 to build a logistic regression based model for classification. We perform 4-fold cross validation in both the stages. We use this technique for both pairwise and multi-classification tasks mentioned in the paper. Further, the technique is used to build both hard-decision and soft-decision based classifiers. While building these classifiers, the number of training samples for each class were equal to each other to account for data imbalance.

### Mutation detection algorithm

We divide the RNA sequence of length *M* into non-overlapping and continuous segments of length *N*. Therefore there are 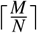segments. For each segment 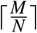, we calculate the transition probability matrix *T^i^*. For the two classes of sequences in question (for example clade 20A and clade 19A mentioned in the paper), we then compare *T^i^* for the sequences in class 1 against the sequences in class 2. We use machine learning steps as stated in the previous section to make this comparison of the transition probability matrices. If the validation accuracy of the classifier built to make this comparison is high (≈0.7 or more), the algorithm claims that there is a set of mutations in the region *i* which make sequences in class 1 distinguishable from sequences in class 2. We do this classification for all the values of *i* and report the set of segments with high validation accuracy as the mutated ones. The transition probability matrix here serves as a transformation from the sequence space into a weighted directed graph where *k*-mers represent the nodes and the weight on an edge (*u, v*) represents the transition probability from node *u* to node *v*. This transformation is robust to insertion, deletion and substitution changes in a way that it aids in the detection of mutations without the need of an alignment step.

## Data availability

The samples are obtained from GISAID and the acknowledgement files with the GISAID sample identifiers are provided on https://github.com/joshuaxiao98/Predicting-the-Emergence-of-SARS-CoV-2-Clades

## Code availability

The code for reproducing the results is provided on https://github.com/joshuaxiao98/Predicting-the-Emergence-of-SARS-CoV-2-Clades

## Author contributions statement

S.J. wrote the manuscript and code for the implementation of the mutation detection, clade classification and clade emergence techniques. X.X. helped with the code development. All authors discussed the results and commented on the manuscript. P.B. and J.B. originated and directed the study.

