## Supplementary Material for "Predicting the Emergence of SARS-CoV-2 Clades"

### ABSTRACT

Evolution is a process of change where mutations in the viral RNA are selected based on their fitness for replication and survival. Given that current phylogenetic analysis of SARS-CoV-2 identifies new viral clades after they exhibit evolutionary selections, one wonders whether we can identify the viral selection and predict the emergence of new viral clades? Inspired by the Kolmogorov complexity concept, we propose a generative complexity (algorithmic) framework capable to analyze the viral RNA sequences by mapping the multiscale nucleotide dependencies onto a state machine, where states represent subsequences of nucleotides and state-transitions probabilities encode the higher order interactions between these states. We apply computational learning and classification techniques to identify the active state-transitions and use those as features in clade classifiers to decipher the transient mutations (still evolving within a clade) and stable mutations (typical to a clade). As opposed to current analysis tools that rely on the edit distance between sequences and require sequence alignment, our method is computationally local, does not require sequence alignment and is robust to random errors (substitution, insertions and deletions). Relying on GISAID viral sequence database, we demonstrate that our method can predict clade emergence, potentially aiding with the design of medications and vaccines.

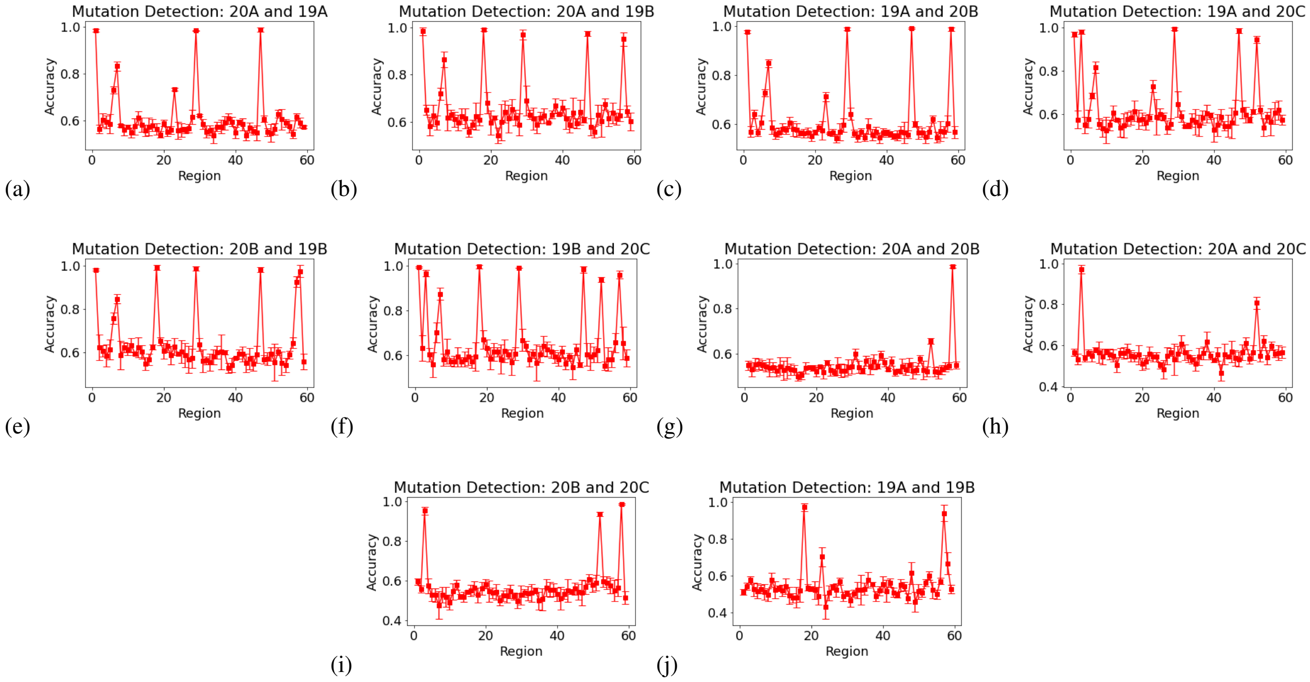


**Figure S1. Mutation detection with region length 500.** SARS-CoV-2 RNA is divided into continuous non-overlapping regions of length 500. Spikes represent the regions where mutations differentiating clade pairs 20A and 19A (a), 20A and 19B (b), 19A and 20B (c), 19A and 20C (d), 20B and 19B (e), 19B and 20C (f), 20A and 20B (g), 20A and 20C (h), 20B and 20C (i), 19A and 19B (j) are detected.

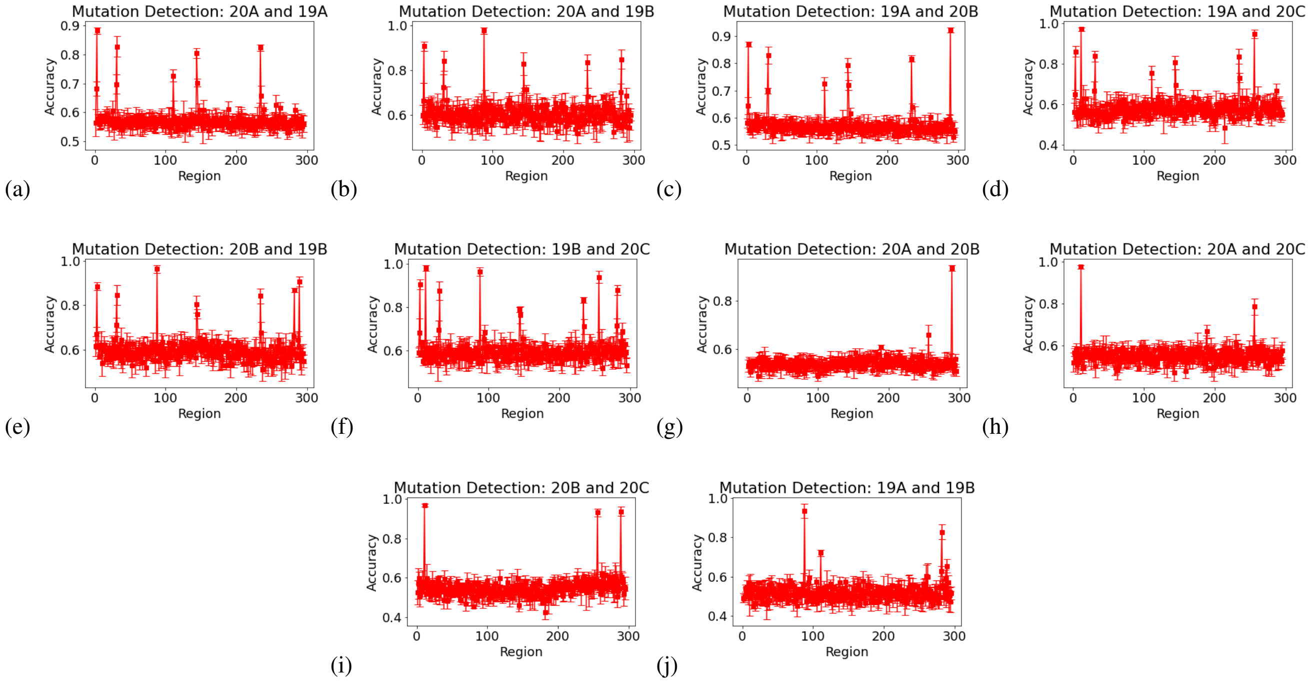


**Figure S2. Mutation detection with region length 100.** SARS-CoV-2 RNA is divided into continuous non-overlapping regions of length 100. Spikes represent the regions where mutations differentiating clade pairs 20A and 19A (a), 20A and 19B (b), 19A and 20B (c), 19A and 20C (d), 20B and 19B (e), 19B and 20C (f), 20A and 20B (g), 20A and 20C (h), 20B and 20C (i), 19A and 19B (j) are detected.

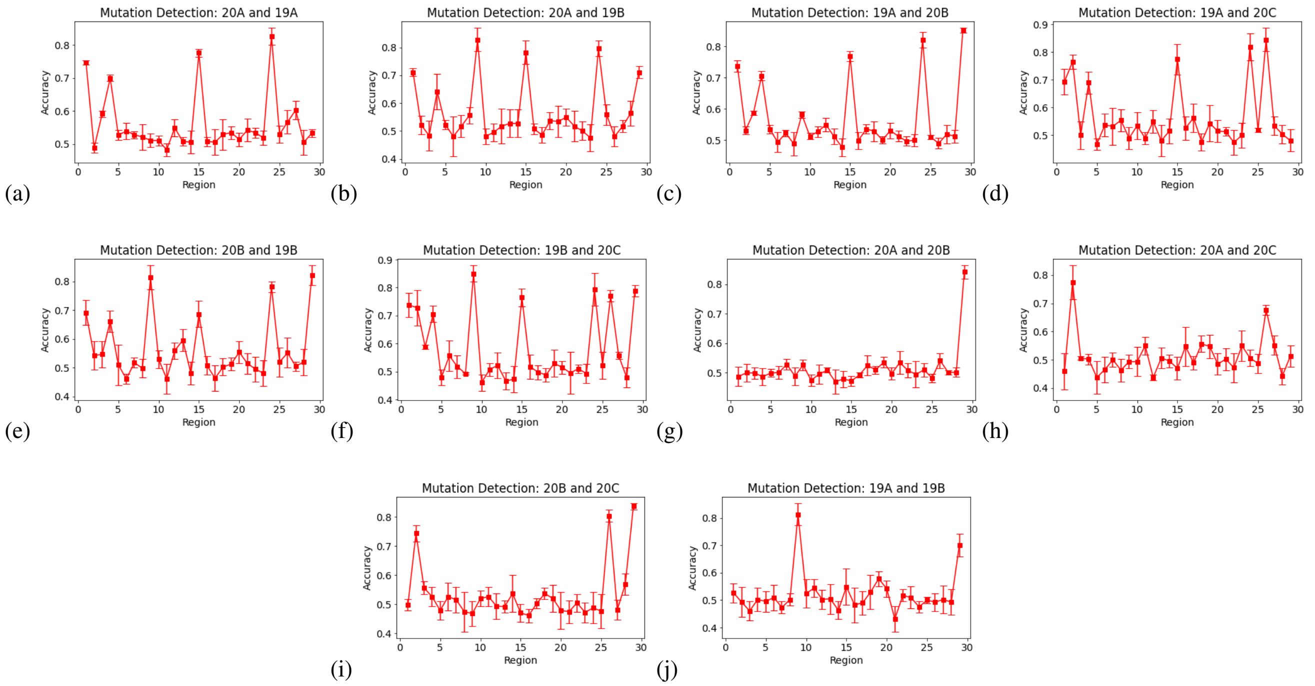


**Figure S3. Mutation detection in the presence of synthetically added noise.** We synthetically added 10% noise in the SARS-CoV-2 viral RNA sequences by adding substitution, insertion and deletion mutations. We show here that the mutations differentiating clades can still be detected, meaning that the detection using state transition probability matrix is robust to noise.

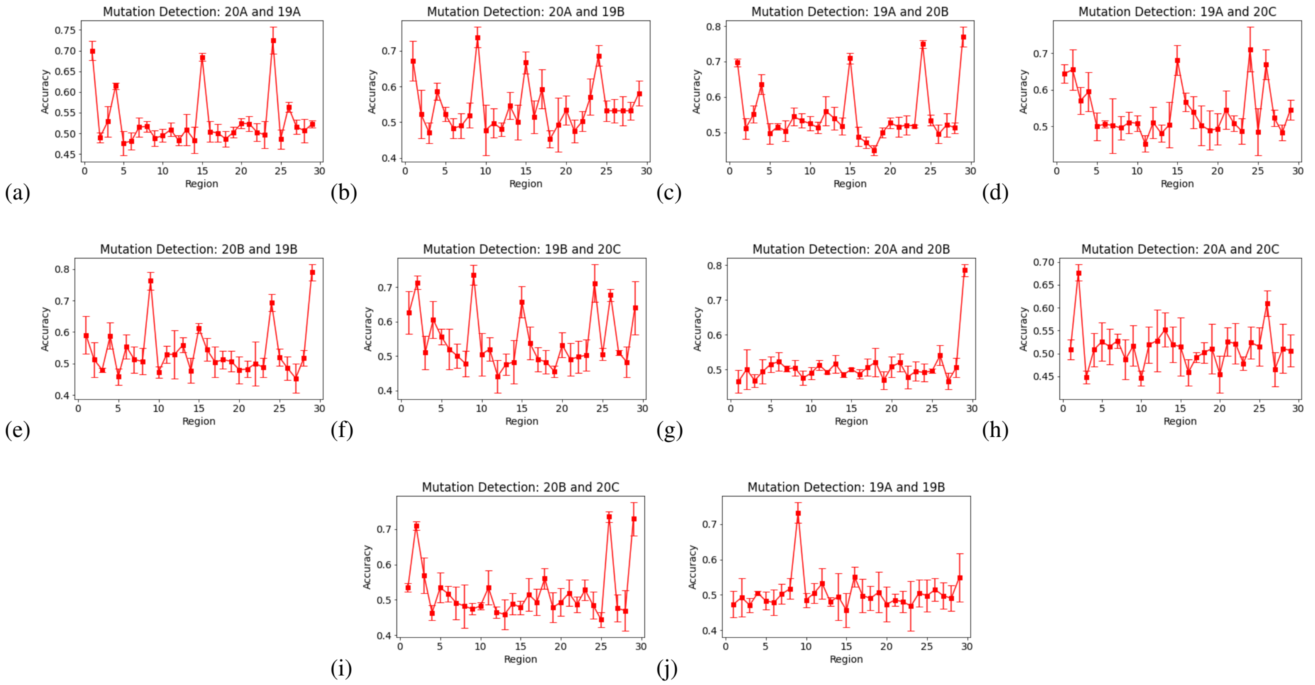


**Figure S4. Mutation detection in the presence of synthetically added noise.** We synthetically added 15% noise in the SARS-CoV-2 viral RNA sequences by adding substitution, insertion and deletion mutations. We show here that the mutations differentiating clades can still be detected, meaning that the detection using state transition probability matrix is robust to noise.

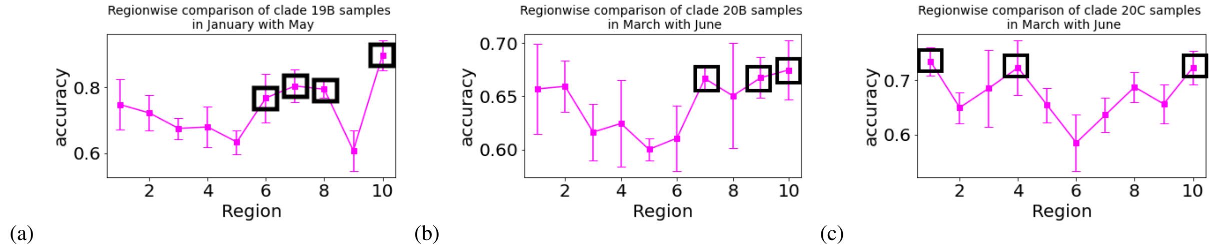


**Figure S5.** **Evolution activity within clades 19B, 20B, 20C.** (a) Regions 6 (15000-18000), 7 (18000-21000), 8 (21000-24000) and 10 (27000-30000) are actively mutating within clade 19B and could correspond to the location of mutations in the new clade that may emerge from 19B in the future. (b) Regions 7 (18000-21000), 9 (24000-27000) and 10 (27000-30000) contributes to maximum classification accuracy and hence are the most actively mutating within clade 20B which indicates the possibility of the emergence of a new clade from 20B with mutations in region 7,9 and 10. (c) Regions 1 (0-3000), 4 (9000-12000) and 10 (27000-30000) are the most actively mutating regions within clade 20C which indicates the possibility of emergence of a new clade from 20C with mutations in these regions.

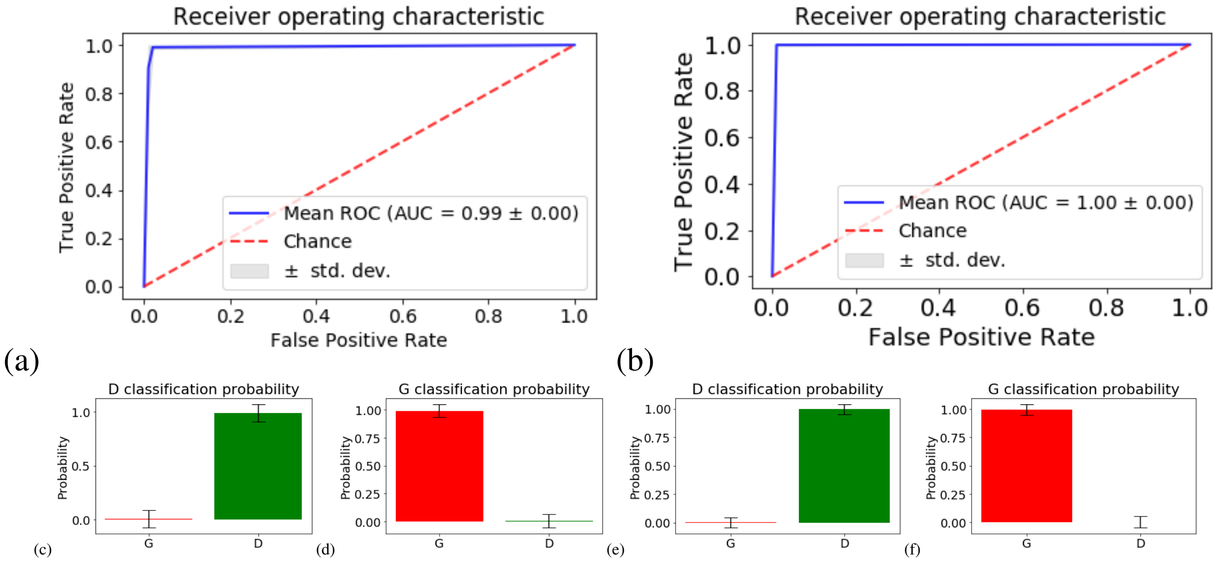


**Figure S6.** **The state machine model can distinguish spike protein mutation sequences from others.** We demonstrate here that the sequences with spike protein mutation (D614G) can be distinguished from others using the state machine transformation. Our state machine model accurately captures the higher order differences and classifies sequences with spike protein mutation from the others. (a), (b) ROC curves for the classifier built using the state transition probability as the feature vector to differentiate between sequences with spike mutation from the others. $k = 4, b = 1$ and $k = 5, b = 1$ for (a) and (b) respectively. (c)-(f) show the classification probability on the validation set for sequences with spike mutation and the others using classifier built with state transition probabilities as features.

**Table S1.** Highly Likely Transitions

| |
|---|
| probability $\geq 0.5$ |
| $aaagc \rightarrow t, aaatc \rightarrow a, aacga \rightarrow a, aactc \rightarrow a, acctc \rightarrow a, actat \rightarrow t, actca \rightarrow a, agcct \rightarrow t, ataac \rightarrow a, atacg \rightarrow t, atcga \rightarrow t, attcg \rightarrow t, atttc \rightarrow a, caacg \rightarrow t, caagc \rightarrow t, cacag \rightarrow a, cagac \rightarrow a, cagcg \rightarrow t, catac \rightarrow a, ccaag \rightarrow a, cccat \rightarrow t, cccca \rightarrow a, cccgc \rightarrow a, ccctc \rightarrow a, ccgag \rightarrow g, ccgcg \rightarrow a, ccggg \rightarrow t, ccggt \rightarrow a, cctcc \rightarrow a, cctgc \rightarrow t, cctgg \rightarrow t, cgaaa \rightarrow t, cgact \rightarrow a, cgatt \rightarrow t, cggac \rightarrow a, cggat \rightarrow g, cgtag \rightarrow t, cgtat \rightarrow a, cgtgg \rightarrow t, cgtgt \rightarrow t, ctaac \rightarrow a, ctacc \rightarrow a, ctagg \rightarrow t, ctcgc \rightarrow a, ctctc \rightarrow a, gaagc \rightarrow t, gaagg \rightarrow t, gaccc \rightarrow t, gacgg \rightarrow t, gactc \rightarrow a, gaggc \rightarrow t, gagtg \rightarrow t, gatgc \rightarrow t, gatgg \rightarrow t, gatta \rightarrow t, gcccg \rightarrow t, gcctc \rightarrow a, gcgag \rightarrow a, gcggc \rightarrow a, gcgtc \rightarrow a, gctac \rightarrow t, gctcc \rightarrow a, gctgc \rightarrow t, gctgg \rightarrow t, ggacc \rightarrow t, ggagc \rightarrow t, ggatc \rightarrow a, ggcag \rightarrow a, gggtg \rightarrow t, gggtt \rightarrow t, ggtga \rightarrow t, ggtgc \rightarrow t, ggtgg \rightarrow t, ggtgt \rightarrow t, gtacc \rightarrow a, gtatc \rightarrow t, gtccc \rightarrow t, gtctc \rightarrow t, gtgcg \rightarrow t, gtggc \rightarrow t, gttag \rightarrow a, gttct \rightarrow t, tactc \rightarrow a, tagtc \rightarrow t, tatcc \rightarrow t, tatcg \rightarrow t, tattc \rightarrow t, tcacc \rightarrow t, tcgag \rightarrow g, tcggt \rightarrow a, tctcg \rightarrow t, tgcag \rightarrow a, tgcga \rightarrow a, tggac \rightarrow a, tggtc \rightarrow a, ttagc \rightarrow t, ttata \rightarrow a, ttatc \rightarrow t, ttccc \rightarrow t, ttcga \rightarrow t, ttcgc \rightarrow a, ttgac \rightarrow a, ttggt \rightarrow g.$ |
| probability $\geq 0.55$ |
| $aacga \rightarrow a, aactc \rightarrow a, agcct \rightarrow t, ataac \rightarrow a, atacg \rightarrow t, attcg \rightarrow t, caagc \rightarrow t, cagcg \rightarrow t, ccaag \rightarrow a, cccat \rightarrow t, cccca \rightarrow a, cccgc \rightarrow a, ccgag \rightarrow g, ccgcg \rightarrow a, ccggg \rightarrow t, ccggt \rightarrow a, cctcc \rightarrow a, cctgc \rightarrow t, cctgg \rightarrow t, cgaaa \rightarrow t, cgact \rightarrow a, cggac \rightarrow a, cggat \rightarrow g, cgtag \rightarrow t, ctaac \rightarrow a, ctacc \rightarrow a, ctcgc \rightarrow a, gaagc \rightarrow t, gaagg \rightarrow t, gaccc \rightarrow t, gaggc \rightarrow t, gcccg \rightarrow t, gcgag \rightarrow a, gcggc \rightarrow a, gcgtc \rightarrow a, gctac \rightarrow t, gctcc \rightarrow a, gctgc \rightarrow t, gctgg \rightarrow t, ggagc \rightarrow t, ggatc \rightarrow a, ggcag \rightarrow a, gggtg \rightarrow t, ggtgg \rightarrow t, ggtgt \rightarrow t, gtacc \rightarrow a, gtatc \rightarrow t, gtccc \rightarrow t, gtctc \rightarrow t, tatcg \rightarrow t, tattc \rightarrow t, tcggt \rightarrow a, tctcg \rightarrow t, tgcga \rightarrow a, tggac \rightarrow a, ttagc \rightarrow t, ttcga \rightarrow t, ttgac \rightarrow a.$ |
| probability $\geq 0.6$ |
| $atacg \rightarrow t, caagc \rightarrow t, ccgag \rightarrow g, ccgcg \rightarrow a, ccggg \rightarrow t, cctcc \rightarrow a, cggac \rightarrow a, cggat \rightarrow g, cgtag \rightarrow t, ctaac \rightarrow a, ctacc \rightarrow a, gaagg \rightarrow t, gaccc \rightarrow t, gaggc \rightarrow t, gcccg \rightarrow t, gcgag \rightarrow a, gctcc \rightarrow a, gctgg \rightarrow t, ggtgt \rightarrow t, gtccc \rightarrow t, gtctc \rightarrow t, tatcg \rightarrow t, ttagc \rightarrow t.$ |
| probability $\geq 0.65$ |
| $ccggg \rightarrow t, cggat \rightarrow g, gctcc \rightarrow a, gtccc \rightarrow t, gtctc \rightarrow t.$ |
| probability $\geq 0.7$ |
| $gtccc \rightarrow t.$ |

**Table S2.** Mutations differentiating different clades (obtained from nextstrain.org)

| Clades | Mutations |
|---|---|
| 19A | Root clade |
| 19B | C8782T, T28144C |
| 20A | C14408T, A23403G |
| 20B | C14408T, A23403G, G28881A, G28882A, G28883C |
| 20C | C14408T, A23403G, C1059T, G25563T |
